## Supplementary Information for "CRISPR prime editing for unconstrained correction of oncogenic *KRAS* variants"

**Table of contents**

Supplementary Figure 1. *KRAS* correction activity of *KRAS*-#1 universal pegRNA in HEK293T/17-*KRAS* library cells.

Supplementary Figure 2. Optimization of pegRNAs to generate *KRAS* heterozygous HEK293T/17 cells.

Supplementary Figure 3. *KRAS* correction activities in *KRAS* heterozygous HEK293T/17 cells.

Supplementary Table 1. Subsitution and indel frequencies at endogenous *KRAS* sites.

Supplementary Table 2. *KRAS*-mutant target sequences used in HEK293T/17-*KRAS* library cells.

Supplementary Table 3. Sequences of gRNA and pegRNAs used in this study.

Supplementary Table 4. Primer sequences used in this study.

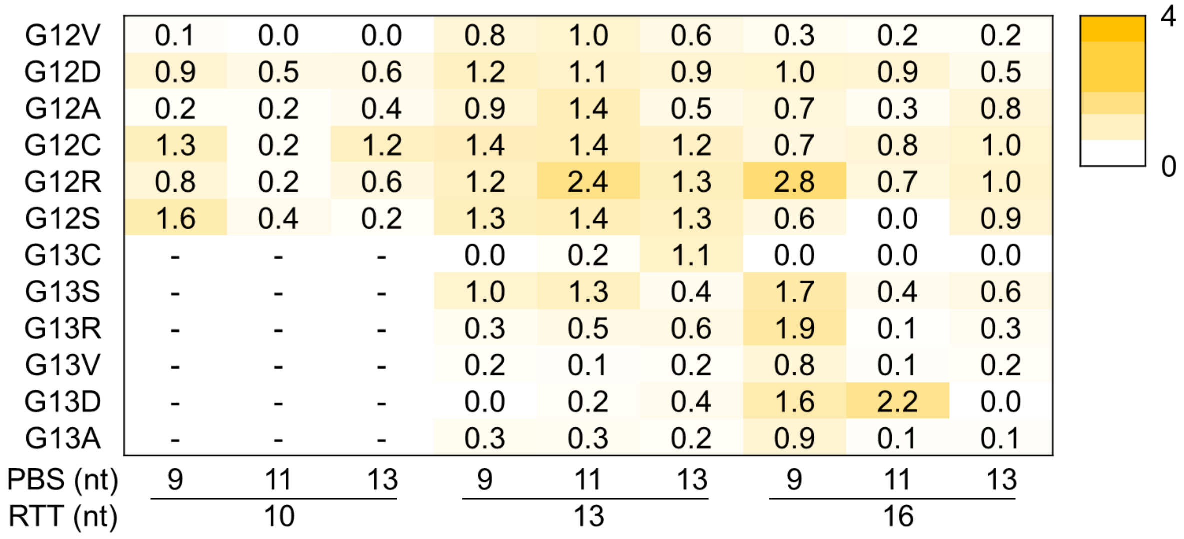


**Supplementary Figure 1**. *KRAS* correction activity of *KRAS*-#1 universal pegRNA in HEK293T/17-KRAS library cells.

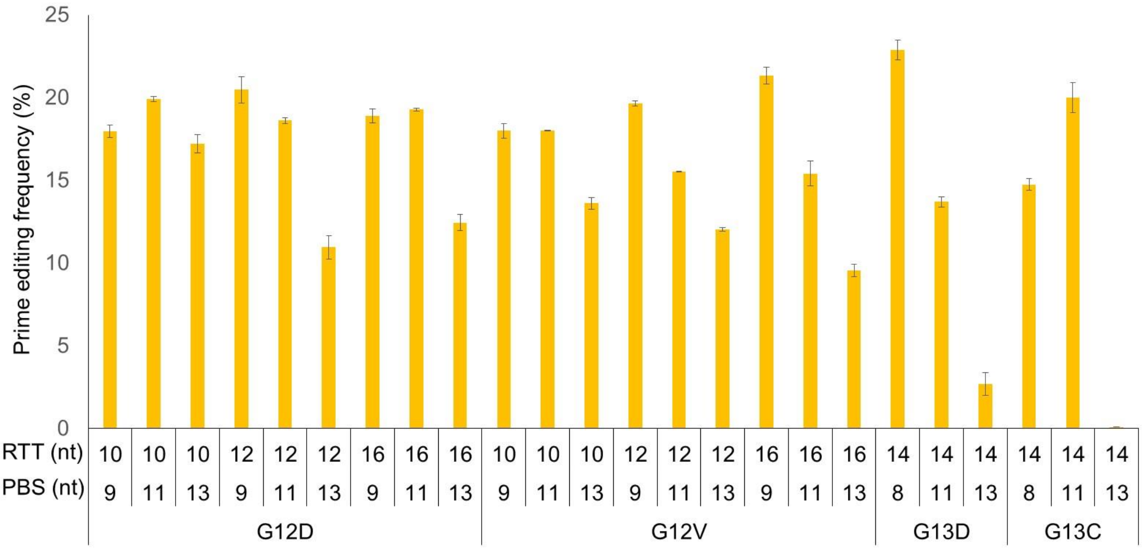


**Supplementary Figure 2**. Optimization of pegRNAs to generate *KRAS* heterozygous HEK293T/17 cells.

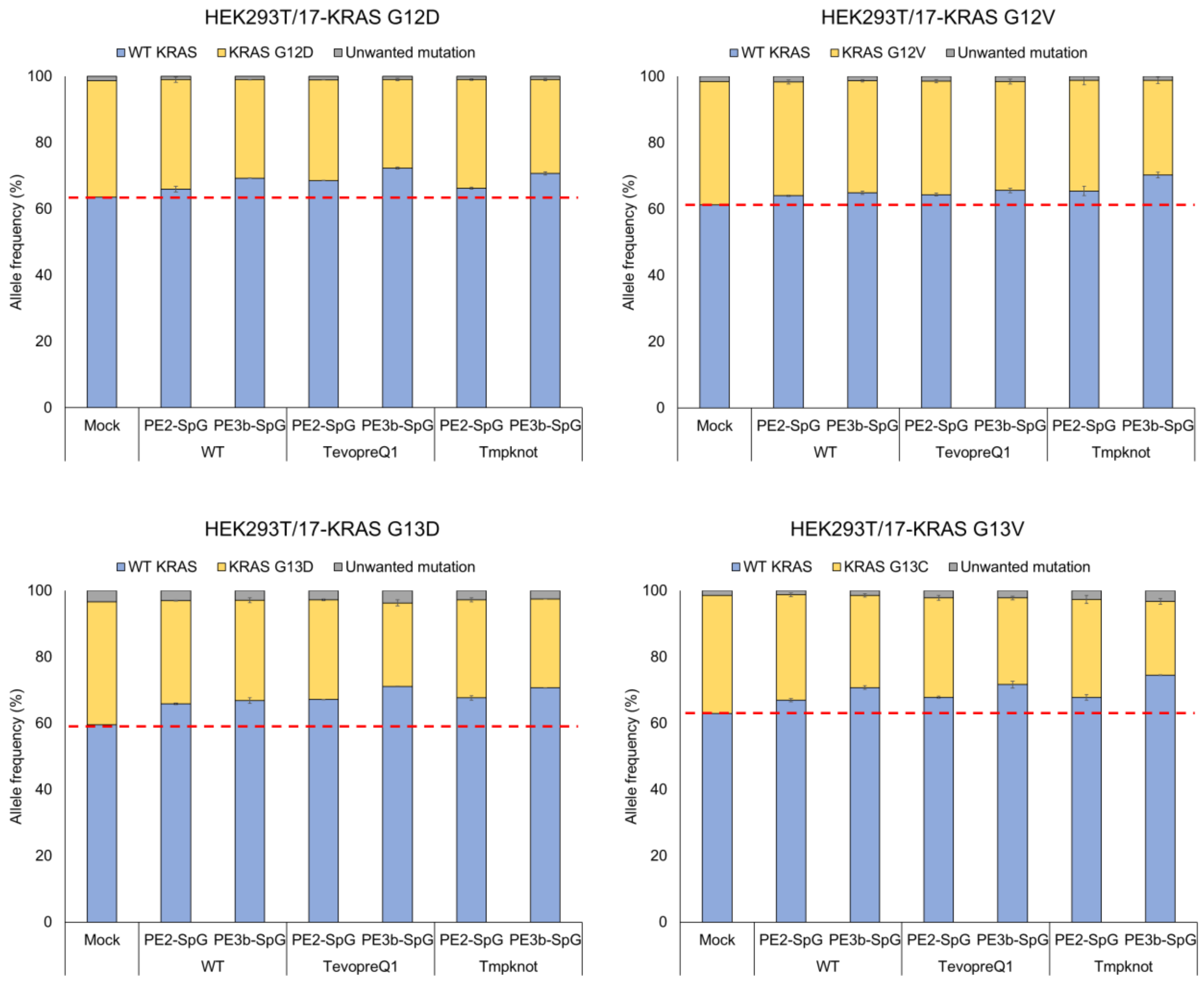


**Supplementary Figure 3**. *KRAS* correction activities in *KRAS* heterozygous HEK293T/17 cells.

**Supplementary Table 1**. Subsitution and indel frequencies at endogenous *KRAS* sites.

| pegRNA names | PBS length (nt) | RT length (nt) | Replicate 1 | | Replicate 2 | |
|---|---|---|---|---|---|---|
| | | | **Substitutions (%)** | **Indels (%)** | **Substitutions (%)** | **Indels (%)** |
| Universal pegRNA #2-1 | 9 | 10 | 1.3% | 0.1% | 1.3% | 0.1% |
| Universal pegRNA #2-2 | 11 | 10 | 1.3% | 0.1% | 1.3% | 0.1% |
| Universal pegRNA #2-3 | 13 | 10 | 1.5% | 0.1% | 1.2% | 0.0% |
| Universal pegRNA #2-4 | 9 | 13 | 1.3% | 0.0% | 1.2% | 0.0% |
| Universal pegRNA #2-5 | 11 | 13 | 1.5% | 0.1% | 1.4% | 0.0% |
| Universal pegRNA #2-6 | 13 | 13 | 1.4% | 0.0% | 1.5% | 0.0% |
| Universal pegRNA #2-7 | 9 | 16 | 1.5% | 0.0% | 1.2% | 0.0% |
| Universal pegRNA #2-8 | 11 | 16 | 1.4% | 0.1% | 1.6% | 0.0% |
| Universal pegRNA #2-9 | 13 | 16 | 1.4% | 0.1% | 1.4% | 0.0% |
| Mock | | | 1.6% | 0.1% | | |

| PE systems | PBS length (nt) | RT length (nt) | RNA scaffolds | Replicate 1 | | Replicate 2 | |
|---|---|---|---|---|---|---|---|
| | | | | **Substitutions (%)** | **Indels (%)** | **Substitutions (%)** | **Indels (%)** |
| PE2-SpG | 13 | 16 | Wildt-type | 0.0% | 0.0% | 0.6% | 0.0% |
| PE3b-SpG | 13 | 16 | Wildt-type | 0.8% | 0.0% | 0.6% | 0.0% |
| PE2-SpG | 13 | 16 | TevopreQ1 | 0.8% | 0.0% | 0.5% | 0.0% |
| PE3b-SpG | 13 | 16 | TevopreQ1 | 0.5% | 0.0% | 0.6% | 0.1% |
| PE2-SpG | 13 | 16 | Tmpknot | 0.6% | 0.2% | 0.6% | 0.1% |
| PE3b-SpG | 13 | 16 | Tmpknot | 0.5% | 0.0% | 0.6% | 0.1% |
| Mock | | | | 0.8% | 0.1% | | |

**Supplementary Table 2**. KRAS-mutant target sequences used in HEK293T/17-KRAS library cells.

| KRAS mutation | KRAS mutant target sequence(5' to 3') | Barcode sequence |
|---|---|---|
| G12V | ATGACTGAATATAAACTTGTGGTAGTTGGAGCTGtTGGCGTAGGCAAGAGTGCCTTGACG | CGAGTAAT |
| G12D | ATGACTGAATATAAACTTGTGGTAGTTGGAGCTGaTGGCGTAGGCAAGAGTGCCTTGACG | TCTCCGGA |
| G12A | ATGACTGAATATAAACTTGTGGTAGTTGGAGCTGcTGGCGTAGGCAAGAGTGCCTTGACG | AATGAGCG |
| G12C | ATGACTGAATATAAACTTGTGGTAGTTGGAGCTtGTGGCGTAGGCAAGAGTGCCTTGACG | GGAATCTC |
| G12R | ATGACTGAATATAAACTTGTGGTAGTTGGAGCTcGTGGCGTAGGCAAGAGTGCCTTGACG | TTCTGAAT |
| G12S | ATGACTGAATATAAACTTGTGGTAGTTGGAGCTaGTGGCGTAGGCAAGAGTGCCTTGACG | ACGAATTC |
| G13C | ATGACTGAATATAAACTTGTGGTAGTTGGAGCTGGTtGCGTAGGCAAGAGTGCCTTGACG | AGCTTCAG |
| G13S | ATGACTGAATATAAACTTGTGGTAGTTGGAGCTGGTaGCGTAGGCAAGAGTGCCTTGACG | GCGCATTA |
| G13R | ATGACTGAATATAAACTTGTGGTAGTTGGAGCTGGTcGCGTAGGCAAGAGTGCCTTGACG | CATAGCCG |
| G13V | ATGACTGAATATAAACTTGTGGTAGTTGGAGCTGGTGtCGTAGGCAAGAGTGCCTTGACG | TTCGCGGA |
| G13D | ATGACTGAATATAAACTTGTGGTAGTTGGAGCTGGTGaCGTAGGCAAGAGTGCCTTGACG | GCGCGAGA |
| G13A | ATGACTGAATATAAACTTGTGGTAGTTGGAGCTGGTGcCGTAGGCAAGAGTGCCTTGACG | CTATCGCT |

**Supplementary Table 3**. Sequences of gRNA for PE3b, pegRNAs and epegRNAs used in this study.

| *KRAS* gRNA or pegRNA name | Target spacer sequence (5' to 3') | PAM | 3' extension sequence (5' to 3') | PBS length (nt) | RTT length (nt) |
|---|---|---|---|---|---|
| G12D mutant pegRNA #1 | AAACTTGTGGTAGTTGGAGC | TGG | GCCAtCAGCTCCAACTACC | 9 | 10 |
| G12D mutant pegRNA #2 | AAACTTGTGGTAGTTGGAGC | TGG | GCCAtCAGCTCCAACTACCAC | 11 | 10 |
| G12D mutant pegRNA #3 | AAACTTGTGGTAGTTGGAGC | TGG | GCCAtCAGCTCCAACTACCACAA | 13 | 10 |
| G12D mutant pegRNA #4 | AAACTTGTGGTAGTTGGAGC | TGG | ACGCCAtCAGCTCCAACTACC | 9 | 12 |
| G12D mutant pegRNA #5 | AAACTTGTGGTAGTTGGAGC | TGG | ACGCCAtCAGCTCCAACTACCAC | 11 | 12 |
| G12D mutant pegRNA #6 | AAACTTGTGGTAGTTGGAGC | TGG | ACGCCAtCAGCTCCAACTACCACAA | 13 | 12 |
| G12D mutant pegRNA #7 | AAACTTGTGGTAGTTGGAGC | TGG | GCCTACGCCAtCAGCTCCAACTACC | 9 | 16 |
| G12D mutant pegRNA #8 | AAACTTGTGGTAGTTGGAGC | TGG | GCCTACGCCAtCAGCTCCAACTACCAC | 11 | 16 |
| G12D mutant pegRNA #9 | AAACTTGTGGTAGTTGGAGC | TGG | GCCTACGCCATCAGCTCCAACTACCACAA | 13 | 16 |
| G12V mutant pegRNA #1 | AAACTTGTGGTAGTTGGAGC | TGG | GCCAaCAGCTCCAACTACC | 9 | 10 |
| G12V mutant pegRNA #2 | AAACTTGTGGTAGTTGGAGC | TGG | GCCAaCAGCTCCAACTACCAC | 11 | 10 |
| G12V mutant pegRNA #3 | AAACTTGTGGTAGTTGGAGC | TGG | GCCAaCAGCTCCAACTACCACAA | 13 | 10 |
| G12V mutant pegRNA #4 | AAACTTGTGGTAGTTGGAGC | TGG | ACGCCAaCAGCTCCAACTACC | 9 | 12 |
| G12V mutant pegRNA #5 | AAACTTGTGGTAGTTGGAGC | TGG | ACGCCAaCAGCTCCAACTACCAC | 11 | 12 |
| G12V mutant pegRNA #6 | AAACTTGTGGTAGTTGGAGC | TGG | ACGCCAaCAGCTCCAACTACCACAA | 13 | 12 |
| G12V mutant pegRNA #7 | AAACTTGTGGTAGTTGGAGC | TGG | GCCTACGCCAaCAGCTCCAACTACC | 9 | 16 |
| G12V mutant pegRNA #8 | AAACTTGTGGTAGTTGGAGC | TGG | GCCTACGCCAaCAGCTCCAACTACCAC | 11 | 16 |
| G12V mutant pegRNA #9 | AAACTTGTGGTAGTTGGAGC | TGG | GCCTACGCCAaCAGCTCCAACTACCACAA | 13 | 16 |
| G13C mutant pegRNA #1 | CTTGTGGTAGTTGGAGCTGG | TGG | TGCCTACGCaACCAGCTCCAAC | 8 | 14 |
| G13C mutant pegRNA #2 | CTTGTGGTAGTTGGAGCTGG | TGG | TGCCTACGCaACCAGCTCCAACTAC | 11 | 14 |
| G13C mutant pegRNA #3 | CTTGTGGTAGTTGGAGCTGG | TGG | TGCCTACGCaACCAGCTCCAACTACCA | 13 | 14 |
| G13D mutant pegRNA #1 | CTTGTGGTAGTTGGAGCTGG | TGG | TGCCTACGtCACCAGCTCCAAC | 8 | 14 |
| G13D mutant pegRNA #2 | CTTGTGGTAGTTGGAGCTGG | TGG | TGCCTACGtCACCAGCTCCAACTAC | 11 | 14 |
| G13D mutant pegRNA #3 | CTTGTGGTAGTTGGAGCTGG | TGG | TGCCTACGtCACCAGCTCCAACTACCA | 13 | 14 |
| Universal pegRNA #1-1 | TATAAACTTGTGGTAGTTGG | AG | AccAGCTCCAACTACCACA | 9 | 10 |
| Universal pegRNA #1-2 | TATAAACTTGTGGTAGTTGG | AG | AccAGCTCCAACTACCACAAG | 11 | 10 |
| Universal pegRNA #1-3 | TATAAACTTGTGGTAGTTGG | AG | AccAGCTCCAACTACCACAAGTT | 13 | 10 |
| Universal pegRNA #1-4 | TATAAACTTGTGGTAGTTGG | AG | GccAccAGCTCCAACTACCACA | 9 | 13 |
| Universal pegRNA #1-5 | TATAAACTTGTGGTAGTTGG | AG | GccAccAGCTCCAACTACCACAAG | 11 | 13 |
| Universal pegRNA #1-6 | TATAAACTTGTGGTAGTTGG | AG | GccAccAGCTCCAACTACCACAAGTT | 13 | 13 |
| Universal pegRNA #1-7 | TATAAACTTGTGGTAGTTGG | AG | TACGccAccAGCTCCAACTACCACA | 9 | 16 |
| Universal pegRNA #1-8 | TATAAACTTGTGGTAGTTGG | AG | TACGccAccAGCTCCAACTACCACAAG | 11 | 16 |
| Universal pegRNA #1-9 | TATAAACTTGTGGTAGTTGG | AG | TACGccAccAGCTCCAACTACCACAAGTT | 13 | 16 |
| Universal pegRNA #2-1 | CGTCAAGGCACTCTTGCCTA | CG | ggTggCGTAGGCAAGAGTG | 9 | 10 |
| Universal pegRNA #2-2 | CGTCAAGGCACTCTTGCCTA | CG | ggTggCGTAGGCAAGAGTGCC | 11 | 10 |
| Universal pegRNA #2-3 | CGTCAAGGCACTCTTGCCTA | CG | ggTggCGTAGGCAAGAGTGCCTT | 13 | 10 |
| Universal pegRNA #2-4 | CGTCAAGGCACTCTTGCCTA | CG | GCTggTggCGTAGGCAAGAGTG | 9 | 13 |
| Universal pegRNA #2-5 | CGTCAAGGCACTCTTGCCTA | CG | GCTggTggCGTAGGCAAGAGTGCC | 11 | 13 |
| Universal pegRNA #2-6 | CGTCAAGGCACTCTTGCCTA | CG | GCTggTggCGTAGGCAAGAGTGCCTT | 13 | 13 |
| Universal pegRNA #2-7 | CGTCAAGGCACTCTTGCCTA | CG | GGAGCTggTggCGTAGGCAAGAGTG | 9 | 16 |
| Universal pegRNA #2-8 | CGTCAAGGCACTCTTGCCTA | CG | GGAGCTggTggCGTAGGCAAGAGTGCC | 11 | 16 |
| Universal pegRNA #2-9 | CGTCAAGGCACTCTTGCCTA | CG | GGAGCTggTggCGTAGGCAAGAGTGCCTT | 13 | 16 |
| PE3b Correction gRNA #1 | GCACTCTTGCCTACGCCACC | AG | N/A | N/A | N/A |
| PE3b Correction gRNA #2 | GTGGTAGTTGGAGCTGGTGG | CG | N/A | N/A | N/A |

| epgRNA name | Target spacer sequence (5' to 3') | PAM | 3' extension sequence (5' to 3') | PBS length (nt) | RTT length (nt) | linker |
|---|---|---|---|---|---|---|
| Universal tevopreQ1 epegRNA #2-9 | CGTCAAGGCACTCTTGCCTA | CG | GGAGCTggTggCGTAGGCAAGAGTGCCTT | 13 | 16 | CTTAAGTA |
| Universal tmpknot epegRNA #2-9 | CGTCAAGGCACTCTTGCCTA | CG | GGAGCTggTggCGTAGGCAAGAGTGCCTT | 13 | 16 | TAATTATA |

**Supplementary Table 4**. Primer sequences used in this study.

| Target sites | | Forward primers | Reverse primers |
|---|---|---|---|
| Endogenous *KRAS* | 1st | CTTAAGCGTCGATGGAGGAG | CCCTGACATACTCCCAAGGA |
| | 2nd | ACACTCTTTCCCTACACGACGCTCTTCCGATCTAGGCCTGCTGAAAATGACTG | GTGACTGGAGTTCAGACGTGTGCTCTTCCGATCTTCATGAAAATGGTCAGAGAAACC |
| Lenti_*KRAS* sequence | 1st | GTGTACGGTGGGAGGCCTAT | GTCCGTCTGCGAGGGTACTA |
| | 2nd | ACACTCTTTCCCTACACGACGCTCTTCCGATCTGCTCGTTTAGTGAACCGTCAG | GTGACTGGAGTTCAGACGTGTGCTCTTCCGATCTACGTGAAGAATGTGCGAGA |
